# Attention to speech: Mapping distributed and selective attention systems

**DOI:** 10.1101/2021.02.13.431098

**Authors:** Galit Agmon, Paz Har-Shai Yahav, Michal Ben-Shachar, Elana Zion Golumbic

## Abstract

Daily life is full of situations where many people converse at the same time. Under these noisy circumstances, individuals can employ different listening strategies to deal with the abundance of sounds around them. In this fMRI study we investigated how applying two different listening strategies – Selective vs. Distributed attention – affects the pattern of neural activity. Specifically, in a simulated ‘cocktail party’ paradigm, we compared brain activation patterns when listeners ***attend selectively*** to only one speaker and ignore all others, versus when they ***distribute their attention*** and attempt to follow two or four speakers at the same time. Results indicate that the two attention types activate a highly overlapping, bilateral fronto-temporal-parietal network of functionally connected regions. This network includes auditory association cortex (bilateral STG/STS) and higher-level regions related to speech processing and attention (bilateral IFG/insula, right MFG, left IPS). Within this network, responses in specific areas were modulated by the type of attention required. Specifically, auditory and speech-processing regions exhibited higher activity during Distributed attention, whereas fronto-parietal regions were activated more strongly during Selective attention. This pattern suggests that a common perceptual-attentional network is engaged when dealing with competing speech-inputs, regardless of the specific task at hand. At the same time, local activity within nodes of this network varies when implementing different listening strategies, reflecting the different cognitive demands they impose. These results nicely demonstrate the system’s flexibility to adapt its internal computations to accommodate different task requirements and listener goals.

**Significance Statement:** Hearing many people talk simultaneously poses substantial challenges for the human perceptual and cognitive systems. We compared neural activity when listeners applied two different listening strategy to deal with these competing inputs: ***attending selectively*** to one speaker vs. ***distributing attention*** among all speakers. A network of functionally connected brain regions, involved in auditory processing, language processing and attentional control was activated when applying both attention types. However, activity within this network was modulated by the type of attention required and the number of competing speakers. These results suggest a common ‘attention to speech’ network, providing the computational infrastructure to deal effectively with multi-speaker input, but with sufficient flexibility to implement different prioritization strategies and to adapt to different listener goals.

## 1. Introduction

Attention is a crucial component of most everyday activities. Given the vast sensory information in our environments and the limited processing resources available, attention is necessary for prioritizing among them (Broadbent, 1958; Posner, 1980; Driver, 2001). However, appropriately allocating processing resources among competing stimuli depends on the specific context and behavioral goals. For example, some circumstances require ***selective attention***, i.e., choosing one information channel over all others. Conversely, in other situations, such as dual-task paradigms or scene-monitoring, multiple channels may be task relevant, requiring ***distributed attention*** to follow them all.

Selective and distributed attention share some common characteristics: They both rely on dynamic interactions between bottom-up sensory processing and top-down executive control, as evident in the co-engagement of sensory cortices and fronto-parietal control networks in both tasks (Corbetta, 1991; Johannsen et al., 1997; Loose et al., 2003; Johnson and Zatorre, 2006; Corbetta et al., 2008; Moisala et al., 2015). However, they differ in the specific cognitive operations they require. To achieve selective attention, one needs to identify the relevant portion of the scene and amplify it, while actively suppressing irrelevant inputs to avoid distraction (Woldorff et al., 1993; Fritz et al., 2007; Zion Golumbic et al., 2013; Fiedler et al., 2019; Hambrook and Tata, 2019). Conversely, to achieve distributed attention, processing resources need to be apportioned among several stimuli, and thus depends critically on the availability of sufficient resources (Lavie et al., 2004, 2014). In the case of speech, distributed attention can be particularly challenging due to inherent linguistic processing bottlenecks (Treisman, 1964; Duncan, 1980; Koelewijn et al., 2014; Bronkhorst, 2015; Kawashima and Sato, 2015; McCloy and Lee, 2015).

Given the similarities and differences in the cognitive operations involved in these two listening strategies, it is interesting to ask whether they engage similar or different neural substrates. Despite an abundance of research regarding the neural mechanisms of selective attention to speech (see review by Miller, 2015), and some work on distributed attention (Getzmann et al., 2016), few studies have directly compared these two types of attention (cf. Tóth et al., 2019; Yuriko Santos Kawata et al., 2020). The goal of this fMRI study was to determine the extent of overlap between the neural substrates functionally involved in selective and distributed attention to concurrent speech, as well as the differences between them.

When asking how the brain deals with competing speech, it is important to consider the nature of the acoustic competition they elicit. A prevalent approach has been to study behavioral or neural responses to brief utterances presented simultaneously from several speakers, i.e. in a fully overlapping manner (Yost et al., 1996; Hugdahl et al., 2000; Jäncke and Shah, 2002; Lipschutz et al., 2002; Gygi and Shafiro, 2012; Scott and McGettigan, 2013). However, this stimulation creates an extreme case of energetic and informational masking, substantially exceeding that encountered under natural condition. Hence, results from such studies might over-emphasize the effects of acoustic masking, rather than attentional effects per se. Here we aimed to study attention to speech under conditions that more closely emulate the natural commotion of hearing multiple speakers. Natural speech is not limited to single utterances but unfolds continuously over time (Hill and Miller, 2010), and its cadence is proposedly important in facilitating attention (Cooke, 2006; Vestergaard et al., 2011; Zion Golumbic et al., 2012; Haegens and Zion Golumbic, 2018; Jones, 2019). Therefore, in this fMRI study, we presented participants with continuous sequences of speech, producing a cacophony of non-stationary, partially overlapping, sounds. In addition, we manipulated the number of concurrent speakers, allowing us to assess the effect of different degrees of acoustic masking. Participants were required to detect a target word spoken by either one pre-determined speaker (Selective Attention) or by any speaker (Distributed Attention). Critically, the acoustic input in both conditions was identical, allowing us to attribute any differences in neural activation during Selective and Distributed attention to the differential cognitive operations they require.

## 2. Materials and Methods

### 2.1. Participants

Thirty-five right-handed Hebrew speaking adults participated in this experiment (16M, 19F; age range: 18.5-35; mean age ±SD: 25 ±4). All participants were right-handed and reported normal hearing, normal or corrected-to-normal vision and no neurological or psychiatric disorders. Prior to the experiment, participants signed a written informed consent in accordance with the declaration of Helsinki. Data from two participants were excluded from the analysis due to incidental findings in their anatomical brain scans (M, 29Y; F, 23Y). One additional participant was excluded from fMRI data analysis due to low performance on the behavioral task, suggesting that she did not follow task instructions (F, 20Y).

### 2.2. Stimuli – attention task

The stimuli and paradigm used here were similar to that used in a previous behavioral study (Lambez et al. 2020). Participants were presented with sequences of short mono-syllabic Hebrew nouns (e.g., *“Kad”;* pitcher in Hebrew). The words were recorded by two male and two female speakers, resulting in four clearly distinguishable streams. We used Audacity (The Audacity Team, 2018) and Matlab (Mathworks, 2012) for audio editing of the individual words, combining them into noun-sequences and equating the volume of each speaker for perceived loudness. Word length varied between 250 ms to 400 ms. Single words from each speaker were concatenated into 12-second long sequences, with inter-stimulus-intervals varying between 500 to 1400 ms. Sequences produced by different speakers were then superimposed, forming either 2-speaker or 4-speaker acoustic scenes, which were presented to both ears (with no spatial cues). The temporal overlap in the 2-speaker condition was carefully controlled such that the audio of the two speakers overlapped for ∼25% of the sequence. In the 4-speaker condition, the temporal overlap was more substantial, with words uttered by 3 different speakers overlapping at any given moment, however common onsets and offsets were avoided. Example stimuli can be downloaded from: https://osf.io/7p84q/?view_only=ae57aeb7c68041f69f14c695df91c87e

### 2.3. Procedure – attention task

The Selective Attention and Distributed Attention conditions were tested in separate runs (Figure 1). In the Selective Attention condition, participants were instructed to attend to one designated speaker and respond to a target word when uttered only by that speaker (2-3 targets per block). A different designated speaker was defined in each Selective Attention run, and participants were familiarized with their voice prior to each run. In order to ensure that participants indeed attended selectively only to the designated speaker, rather than follow all the speakers, Selective Attention blocks also contained 1-2 ‘catch-stimuli’, where the target word was spoken by one of the task-irrelevant speakers. If participants performed false-alarms in response to these catch-stimuli, this would indicate they did not correctly follow task-instructions. Conversely, in the Distributed Attention condition, participants were required to respond to the target word spoken by <u>**any**</u> of the speakers (2-3 targets per block). Subjects responded to the target word by pressing a button using their left index finger, to minimize motor activation adjacent to language areas in the left hemisphere. The same target word (*“Etz”*, a tree in Hebrew) was used in both Selective-and Distributed-Attention conditions.

**Figure 1.**
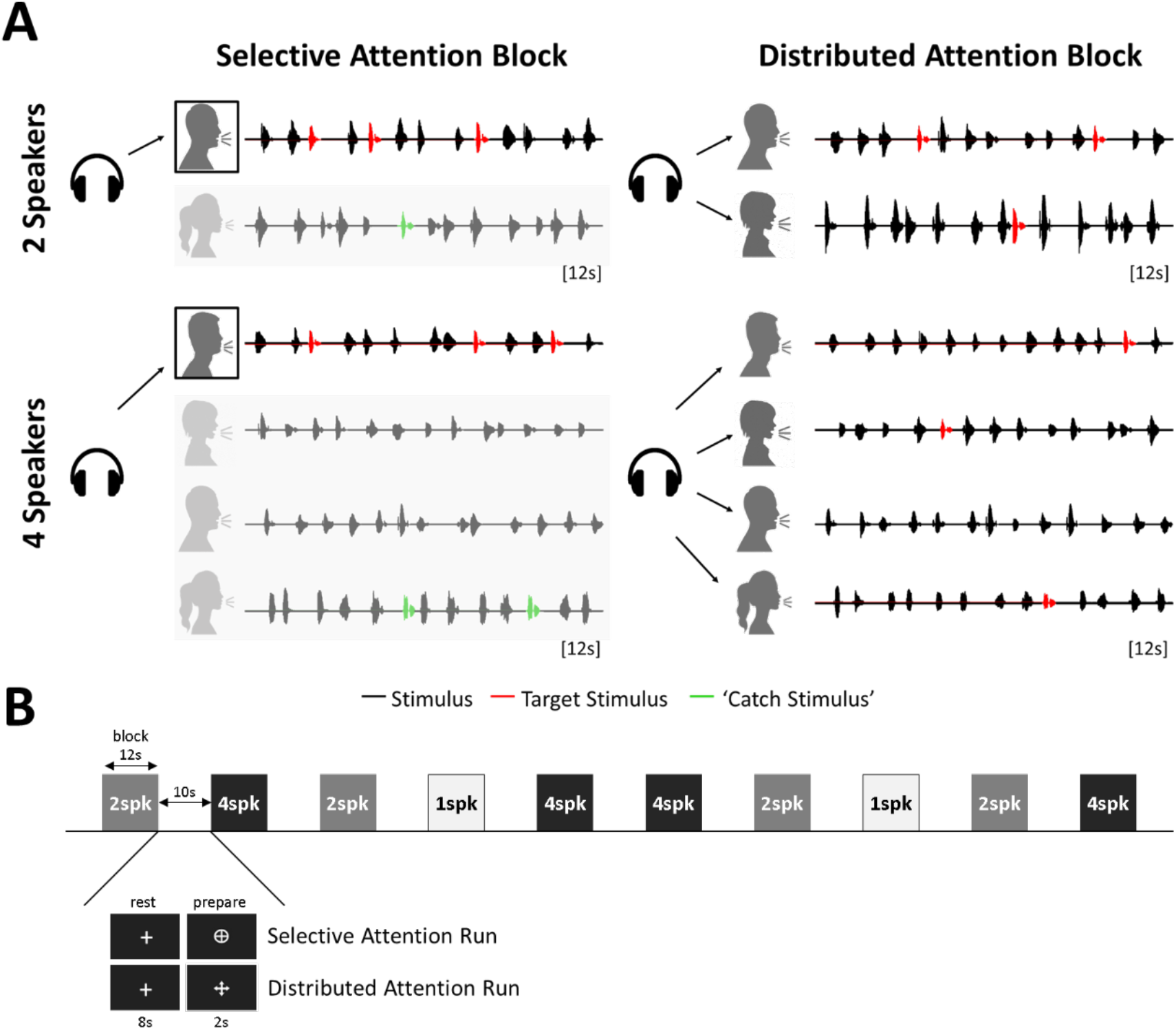
Experimental design and task. (A) Illustration of the stimuli presented in the 2-speaker and 4-speaker blocks, separately for the Selective and Distributed conditions. Stimuli were lists of mono-syllabic nouns (black), with one word serving as a target (red). In Selective Attention blocks (left) participants were required to attend to one designated speaker and respond to target words uttered only by this speaker (red). Utterances of the target by one of the other speakers are considered ‘catch-stimuli’ (green) and should be ignored. In Distributed Attention blocks (right) participants were required to respond to a target word uttered by <u>any</u> of the speakers (red). (B) Illustration of the block-design structure within each run. The number of concurrent speakers was manipulated between blocks (1spk, 2spk, and 4spk represent blocks of 1 speaker, 2 speakers and 4 speakers, respectively). Different visual icons were presented prior to and throughout each run, to instruct and remind participants if they should perform the Selective or Distributed attention task in that particular run.

The experiment consisted of six runs, with three runs per attentional condition (Selective/Distributed attention), presented in pseudo-random order. Each run consisted of ten blocks (Figure 1B), with the Number of Speakers manipulated across blocks (four blocks with 2 speakers; four blocks with 4 speakers and two blocks with a single speaker, serving as a baseline). Experimental blocks were 12-second long, and were interleaved with 10-second long rest blocks (8-second fixation; 2-second preparation icon). Two distinct icons were used to visually indicate which attentional task was to be performed in each run: a circled-cross for the Selective Attention condition and a cross with outward arrows for the Distributed Attention condition (see Figure 1B) The icons appeared 2-seconds before the start of each experimental block and remained on the screen throughout the block to serve as a reminder. In total, each run lasted approximately 4 minutes and the duration of the full attention experiment was 24 minutes.

### 2.4. Procedure – speech localizer task

To isolate auditory and speech-sensitive cortical regions, we used a speech localizer task (Stoppelman et al., 2013). The localizer was implemented in a block design, with 6 experimental blocks interleaved with rest blocks. Experimental blocks included 3 blocks of listening to speech (Speech) and 3 blocks of listening to signal correlated noise (SCN), a well-matched auditory baseline (Rodd et al. 2005, Davis et al. 2007). The Speech>SCN contrast was previously shown to effectively delineate speech and language areas from primary and associative auditory areas. Speech and SCN blocks, 15-second long each, were presented in a random order that was constant across subjects and rest periods of 12.5-seconds were interleaved between the experimental blocks. Participants performed a simple target detection task, pressing a button whenever they heard “blip” sounds randomly placed throughout the entire experiment. For more information about the stimuli and procedure see (Stoppelman et al., 2013).

### 2.5. Scanning parameters

MRI data were collected using a Siemens 3T MAGNETOM Prisma scanner (Siemens Medical Systems, Erlangen, Germany), located at the Alfredo Federico Strauss Center for Computational Neuro-imaging at Tel-Aviv University (https://mri.tau.ac.il/). Scanning was conducted using a 64-channel head-coil. Behavioral responses during fMRI scans were recorded with a response box (932 interface unit, Current Designs, Philadelphia, PA, USA), placed on the participant’s lap under the fingers of their left hand. Auditory stimuli were presented using MRI compatible in-ear headphones (S14, Sensimetrics, Gloucester, MA, USA). Head motion was minimized by padding placed around the head and by instructing the subjects to stay still throughout the entire scan.

T1-weighted anatomical scans were acquired using an MP-RAGE protocol (TR=2530 ms, TE=2.45 ms, flip angle=7°, FOV=224×224 mm^2^, matrix size: 224×224, 176 1-mm thick slices, no gap, resulting in 1×1×1 mm^3^ voxel size, parallel imaging using GRAPPA 2). The scan lasted about 4 minutes.

T2*-weighted functional scans during the Attention task were acquired using an EPI sequence (TR=2000 ms, TE=30 ms, flip angle=80°, FOV=180×208 mm^2^, matrix=90×104, 64 2-mm thick slices, no gap, resulting in a voxel size of 2×2×2 mm^3^, interleaved acquisition, with a multi-band acceleration factor of 2). For each subject, a total of 714 functional volumes were acquired in the attention experiment, divided into 6 runs, such that each run consisted of 119 volumes. For the speech localizer, 95 volumes were acquired in a single run, using the same EPI sequence with a longer TR (TR=2500 ms). This adjustment was necessary in order to synchronize the total block length with a round number of acquisition volumes.

### 2.6 Data analysis – behavioral responses

In the Attention task, true detection (a *hit*) was defined as a button press issued between 300 and 1800 ms after the target word. Any button press outside this critical time window was deemed as a *false alarm*. Repeated button presses after the first one issued in the critical time window were also tagged as false alarms. If no button press was detected within the critical time window, the target was tagged as a *miss*. Behavioral performance was used to screen the data, such that participants with low hit rates were excluded from the analysis (beyond two standard deviations from the sample mean).

To investigate differences in accuracy and reaction times (RTs) between the conditions, we performed linear mixed effects regression analysis using R’ lme4 package (Bates et al., 2015). For RTs we used the logarithmic transformation of reaction times to hits only. For accuracy we used a generalized (logistic) linear mixed effects regression model using all the hits and misses. Mixed effects regression analysis was chosen in order to account for between-subject variability and the correlational structure of the effects of interest (Baayen et al., 2008). In addition, logistic mixed effect regression is the most accurate way to parametrically analyze binomial data, such as performance in detection tasks, in repeated-measures designs (Jaeger, 2008). Both the accuracy and RT models included random intercepts and random Attention Type slopes, adjusted by subjects. Attention Type was sum coded (i.e., the coefficient represents the difference between the Distributed Attention condition and the grand mean), and Number of Speakers was forward-Helmert-coded (i.e., one coefficient represents the difference between the 1-speaker condition and the average of 4-and 2-speaker conditions, and the second coefficient represents the difference between the 2-speaker condition and the 4-speaker condition). The p-values for the RT model were estimated based on Satterthwaite approximation of degrees of freedom (Satterthwaite, 1946), implemented in R’ lmerTest package (Kuznetsova et al., 2017). For the hit rate model, p-values were based on asymptotic Wald tests which are included in the summary of R’ glmer function.

### 2.7. Data analysis – functional data

#### 2.7.1. Preprocessing

Data analysis was performed using AFNI (Cox, 1996). We removed the first four TRs of each functional run, to allow the signal to reach equilibrium. All volumes were time-shifted and registered to a reference volume (10^th^ volume after the T1-weighted scan of the Attention task), motion-corrected, scaled to percent-signal-change from the mean, temporally de-spiked and spatially smoothed to reach a 6 mm FWHM Gaussian kernel (using AFNI’ function 3dBlurToFWHM). All functional volumes with motion of >0.9 or >0.9° were excluded from the analysis (0.1% of all Attention task volumes, 1% for the subject with the most movement). The T1-weighted images were corrected for signal inhomogeneity, skull-stripped, and aligned to the same reference volume as the functional images.

#### 2.7.2. Individual-level analysis – attention task

For each participant, we fitted a general linear model (GLM) via AFNI’ program 3dDeconvolve. The model included five regressors for the conditions of interest: Selective 2-speakers, Selective 4-speakers, Distributed 2-speakers, Distributed 4-speakers and 1-speaker condition (each regressor was estimated based on 12 blocks).

Additional regressors were included for the three elements of a quadratic polynomial in each of the six runs (a total of 18 additional regressors), to account for signal of no-interest (e.g., due to drift or cardiac/respiratory cycles). The degree of the polynomial was the highest possible given run duration, based on 3dDeconvolve’ automatic option of polynomial degree. In addition, the model included six regressors to account for motion (x, y, z, roll, pitch, and yaw), yielding a total of 29 regressors in the GLM. The hemodynamic response function assumed by the model was AFNI’ BLOCK function, convolved with the duration of the experimental block (12 seconds).

#### 2.7.3. Group-level analysis – attention task

For the group-level analysis, we first converted all of the GLM results from the individual-level analysis into an MNI template (MNI152). We ran two main analyses. First, to determine which brain areas were activated in each attention task, we looked at the contrast between conditions with multiple speakers (average of 2-speakers and 4-speakers) vs. 1-speaker. This analysis was performed for each Attention Type separately (Selective/Distributed), yielding statistical parametric maps for each task, that allowed us also to assess the degree of overlap between them.

The union of these two activation maps was then used as a mask for a follow up 2×2 repeated-measures ANOVA analysis directly comparing the effects of Attention Type (Selective/Distributed) and Number of Speakers (2-speakers/4-speakers).

All statistical analyses were corrected for multiple comparisons using AFNI’ 3dClustSim function (100000 iterations), that runs simulations of noise-only activation in order to obtain a data-driven threshold for statistical significance at a cluster-level corrected p<0.003. The uncorrected threshold required to achieve this cluster-corrected p-value was p=0.01 for the Number of Speakers contrast, and p=0.05 for the Distributed vs. Selective contrast and for the interaction.

#### 2.7.4. Individual-level and group-level analyses – Speech Localizer task

Preprocessing of the speech localizer task was done similarly to the preprocessing of the Attention task (see *Preprocessing*). Then, for each participant, we fitted a GLM to the preprocessed data, using AFNI’ 3dDeconvolve. The GLM included one regressor for the Speech condition and one regressor for the SCN condition, in addition to a quadratic polynomial and six movement parameters (10 regressors in total). AFNI’ BLOCK function was convolved with a 15 seconds block to construct the design matrix.

Next, we performed a group-level analysis, to generate a map of speech processing areas. We first converted all the GLM results from the individual-level analysis into an MNI template (MNI 152). We compared Speech with SCN by calculating a t-test (AFNI’ ttest++), and corrected for multiple comparisons using 3dClustSim to reach a corrected p-value of 0.003 (uncorrected threshold per voxel p=0.00001). We also delineated auditory areas via a group level t-test comparing SCN with rest, using the same statistical thresholds. For visualization purposes, we overlaid the contour of these two maps on the results of the multiple vs. single speaker contrast, to assess the prevalence of attention-related effects within and outside speech areas.

#### 2.7.5. Functional connectivity analysis

In this analysis, we examined how functional connectivity changed during the task compared to the rest blocks, and under the different attention conditions. To limit the complexity of this analysis, we focused on a subset of regions of interest (ROIs) where there were significant simple effects of Attention Type or Number of Speakers (see Table 5). Pairwise correlations between ROIs were computed on the GLM residuals of the individual-level analysis. The residuals capture the variability that remains after the effects of task and movement are accounted for, yet may still include task-relevant variability that is independent of BOLD amplitude (e.g.,Tran et al., 2018).

**Table 5.** ROIs used for the functional connectivity analysis

| Region | # voxels | Size (cm<sup>3</sup>) | Center of Mass (MNI coordinates) |
| --- | --- | --- | --- |
| Left STG | 1751 | 14.0 | -55, -23, 6 |
| Right STG | 1417 | 11.3 | 58, -17, 5 |
| Right insula lobe | 1113 | 8.9 | 40, 20, 2 |
| Left IPS | 863 | 6.9 | -35, -45, 52 |
| Left insula lobe | 858 | 6.9 | -36, 19, 2 |
| Right MFG | 828 | 6.6 | 41, 21, 35 |
| Right IPS | 199 | 1.6 | 37, -51, 47 |

STG, superior temporal gyrus; IPS, inferior parietal lobule; MFG, middle frontal gyrus.

To this end, we spliced and concatenated the residual time-series acquired in each condition-combination of Attention Type and Number of Speakers. To assess non-task resting-state functional connectivity we concatenated data from the Rest periods (Fair et al., 2007), acquired separately during Distributed/Selective runs. To reduce the spillover from experimental blocks to Rest TRs, we removed two TRs from the beginning of each Rest period before concatenation.

Connectivity strength was calculated between all ROI-pairs (within hemisphere), by calculating the Pearson’s correlation between their averaged residual time-series (AFNI’ 3dNetCorr program (Taylor and Saad, 2013). This was performed separately for each subject and condition (including Rest). Subjects’r-coefficients were Fisher-z transformed, and these scores were used as the dependent variable in a linear mixed effects regression, testing whether connectivity strength between ROI-pairs was affected by condition (R’ lme4; (Bates et al., 2015). The regression model included Attention Type, Number of Speakers and their interaction as fixed factors, and by-subject random intercepts and slopes. Attention Type was sum coded (i.e. coefficient represents the difference between the Distributed Attention condition and the grand mean). Number of Speakers was forward-Helmert-coded: one coefficient represents the difference between Rest and the average of all other conditions (1-, 2-and 4-speakers); the second coefficient represents the difference between the 1-speaker condition and the average of 4-and 2-speaker conditions; and the third coefficient represents the difference between the 2-speaker condition and the 4-speaker condition.

## 3. Results

### 3.1. Behavioral results

Two participants were excluded from all analyses due to incidental findings in their MRI scans. One additional subject was excluded due to low hit rate (59%) and high percentage of false alarms (41%). Four participants exhibited poor performance in only a single run (hit rate <50%). These participants were included in the sample, but the low-performance run was removed. Thus, behavioral and functional results are reported on a sample of 32 participants.

Mean hit rates were high across all conditions (90±5%) with few false alarms (6±4%). Linear mixed effects regression indicated a significant effect of Number of Speakers on hit-rates, with reduced accuracy as the number of speakers increased (p<0.0001 in both the 1-speaker vs. multiple speakers, and 2-speaker vs. 4-speaker contrasts). There was also an effect of Attention Type on hit-rates (p=0.002), indicating slightly higher hit rates in the Distributed vs. Selective Attention condition, possibly because of the presence of catch-stimuli in the latter condition (Figure 2A, Table 1). The analysis of reaction times (RTs) yielded similar main effects for the Number of Speakers contrasts, with longer RTs as the number of speakers increased (p<0.0001 in both the 1-speaker vs. multiple speakers, and 2-speaker vs. 4-speaker contrasts), but no effect of Attention Type and no significant interactions on RT were observed (Figure 2B, Table 2). The proportion of false alarms did not differ significantly between conditions, indicating good compliance with the two tasks [Distributed Attention 6.9±3.3%; Selective Attention 5.6±4.7%; paired t-test: t(31)=1.1, p=0.14].

**Figure 2.**
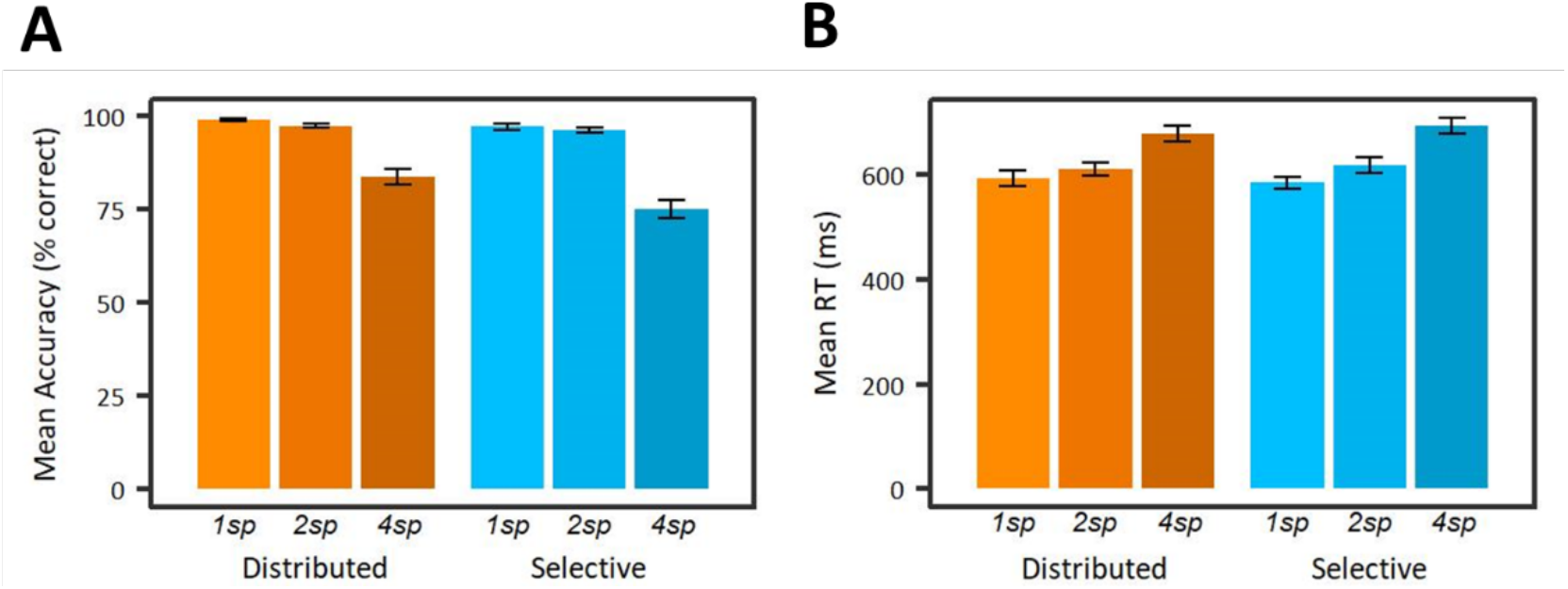
(A) Mean accuracy (hit rate) and (B) Reaction times averaged across participants and runs, in the Distributed (orange) and Selective (blue) Attention conditions, as a function of Number of Speakers (encoded by color-shade). Error bars indicate standard deviations across participants. 1sp = 1-speaker; 2sp = 2-speakers; 4sp = 4-speakers

**Table 1.** Summary of the RT model (RTs in ms were log-transformed).

| Contrast | $\beta \pm \text{SD}$ | t-statistic | p-value |
| --- | --- | --- | --- |
| Attention (Distributed vs. grand mean) | -1.8±3.5 | -0.5 | 0.6 |
| Number of Speakers (1 vs. multiple) | -60.3±5.5 | -11 | <0.0001 |
| Number of Speakers (2 vs. 4) | -70±5 | -13.6 | <0.0001 |
| Attention×Number of Speakers (1 vs. multiple) | 10±5.5 | 1.8 | 0.07 |
| Attention×Number of Speakers (2 vs. 4) | 3.9±5.1 | 0.8 | 0.4 |

**Table 2.** Summary of the hit rate model.

| Contrast | $\beta \pm \text{SD (log-ms)}$ | z-statistic | p-value |
| --- | --- | --- | --- |
| Attention (Distributed vs. grand mean) | 0.4±0.1 | 3.1 | 0.002 |
| Number of Speakers (1 vs. multiple) | 1.8±0.3 | 6 | <0.0001 |
| Number of Speakers (2 vs. 4) | 2.2±0.1 | 14.6 | <0.0001 |
| Attention×Number of Speakers (1 vs. multiple) | 0.4±0.3 | 1.2 | 0.2 |
| Attention×Number of Speakers (2 vs. 4) | -0.1±0.1 | -0.7 | 0.5 |

### 3.2. Whole brain group-analysis results

Figure 3 and Table 3 show the results of a whole-brain group-level analysis indicating which brain regions were significantly more active in the multi-speaker conditions relative to the single-speaker condition (multiple speakers > 1-speaker contrast). This analysis was performed separately for the Selective and Distributed conditions (cyan and orange, respectively). Remarkably, there was substantial overlap between the activation maps obtained for the two Attention Types (overlapping regions are shown in purple; 81.5% of the voxels activated in the Distributed Attention condition were also activated by the Selective Attention condition). The overlapping regions included bilateral responses along the superior temporal gyrus (STG) and the superior temporal sulcus (STS), the insulas, inferior frontal gyrus (IFG) and supplementary motor area (SMA), as well as some unilateral responses in left middle frontal gyrus (MFG) and right cingulate cortex. Notably, the bilateral clusters of activation in STG/STS were not limited to the auditory cortices (gray outline; based on an independent auditory localizer) but extended both posteriorly and anteriorly, overlapping with regions associated with speech processing (black outline; based on an independent speech localizer). Alongside the considerable overlap in regions active in both attention conditions, a few regions were found to be active only in the Selective Attention condition. These included the right MFG, left inferior parietal sulcus (IPS) and bilateral precuneus. Interestingly, we found no regions that were active only in the Distributed but not in the Selective attention condition.

**Figure 3.**
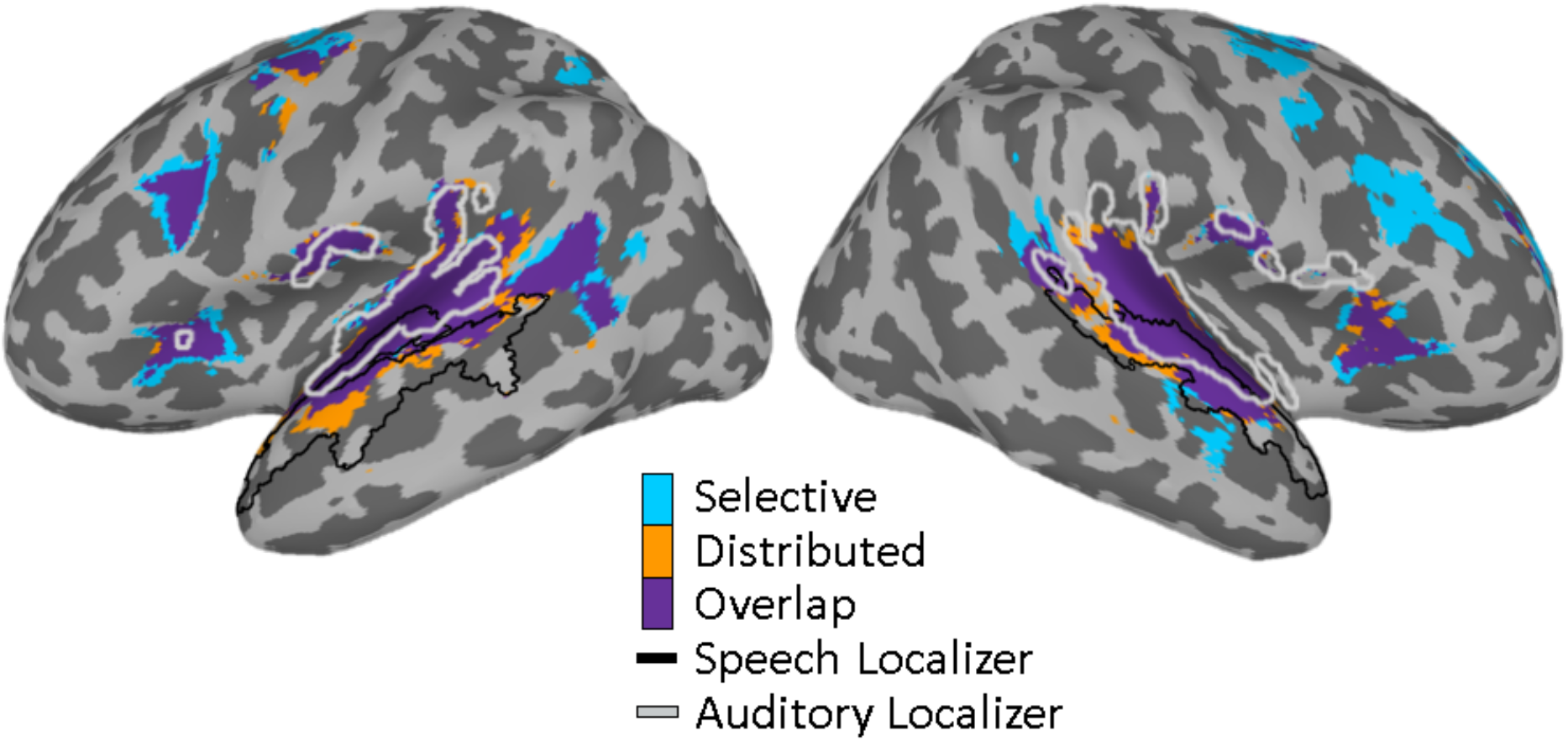
Regions showing enhanced responses to the multiple speakers vs. single speaker condition (p<0.003, corrected). Results are shown separately for Selective (cyan) and Distributed attention (orange), as well as for the overlap between them (purple). Black and gray contours denote results from the auditory and speech localizer task, indicating the regions that responded to signal-correlated-noise (SCN) vs. rest (gray) and regions that responded to speech vs. SCN (black, all ps<0.003, corrected). Results are projected on an inflated brain (MNI152 template, AFNI).

**Table 3.** Significant clusters showing enhanced responses to multiple speakers vs. a single speaker, separately for the Selective and Distributed attention conditions

| (a) Distributed Attention (multiple speakers > 1 speaker) |  |  |  |  |
| --- | --- | --- | --- | --- |
| Region | # voxels | Size (cm<sup>3</sup>) | Peak (MNI coordinates) | t-statistic (sample mean) |
| Left STG | 2993 | 24.0 | -50, -14, 4 | 5.7 |
| Right STG | 2385 | 19.1 | 66, -18, 6 | 6.0 |
| Right SMA | 417 | 3.3 | 4, 10, 60 | 4.2 |
| Right Caudate Nucleus | 225 | 1.8 | 26, 26, 8 | 4.1 |
| Left Precentral Gyrus | 192 | 1.5 | -44, -4, 50 | 4.5 |
| Left Insula | 166 | 1.3 | -30, 24, 8 | 4.6 |
| (b) Selective Attention (multiple speakers > 1 speaker) |  |  |  |  |
| Region | # voxels | Size (cm<sup>3</sup>) | Peak (MNI coordinates) | t-statistics (sample mean) |
| Left STG | 2509 | 20.1 | -48, -14, 2 | 5.5 |
| Right STG | 1882 | 15.1 | 64, -18, 6 | 5.6 |
| Right Middle Cingulate Cortex | 450 | 3.6 | 10, 26, 40 | 4.1 |
| Left Precentral Gyrus | 390 | 3.1 | -40, -6, 48 | 4.3 |
| Left Insula | 249 | 2.0 | -32, 18, 6 | 4.4 |
| Right SMA | 236 | 1.9 | 16, 4, 66 | 4.1 |
| Left IFG | 229 | 1.8 | -46, 18, 26 | 4.4 |
| Right Caudate Nucleus | 162 | 1.3 | 24, 22, 8 | 4.1 |
| Right Precuneus | 83 | 0.7 | 8, -56, 50 | 3.9 |
| Left Superior Parietal Lobule | 53 | 0.4 | -24, -52, 46 | 4.2 |

STG, superior temporal gyrus; SMA, supplementary motor area; IFG, inferior frontal gyrus.

We next assessed how Attention Type and Number of Speakers affected the level of neural activation within the regions identified in Figure 3, using a 2×2 ANOVA (full results listed in Table 4). As shown in Figure 4A, the main effect of Attention Type revealed that responses in bilateral clusters along the STS/STG were stronger when performing Distributed vs. Selective Attention. Conversely, responses in the right MFG and left IPS were enhanced during Selective Attention. The results of this ANOVA further showed a significant main effect of Number of Speakers in a large cluster along bilateral STS/STG as well as bilateral insulas (Figure 4B), where responses were stronger in the 4-speaker vs. 2-speaker conditions. Additionally, there was a significant interaction between Attention Type and Number of Speakers in bilateral insulas and in the right MFG (Figure 4C), where the effect of Number of Speakers was stronger for the Distributed Attention condition compared to the Selective Attention condition. The fMRI data is available at: https://osf.io/7p84q/?view_only=ae57aeb7c68041f69f14c695df91c87e.

**Table 4.** Significant clusters detected in the 2×2 ANOVA (p<0.003, corrected).

| (a) Number of Speakers (4>2) |  |  |  |  |
| --- | --- | --- | --- | --- |
| Region | # voxels | Size (cm<sup>3</sup>) | Peak (MNI coordinates) | t-statistic (sample mean) |
| Left STG | 2601 | 20.8 | -44, -28, 10 | 4.6 |
| Right STG | 2301 | 18.4 | 66, -20, 6 | 4.8 |
| Right Middle Cingulate Cortex | 1632 | 13.1 | 8, 26, 36 | 4.4 |
| Right insula lobe | 731 | 5.8 | 38, 22, -2 | 6.7 |
| Left insula lobe | 543 | 4.3 | -30, 24, 8 | 5.9 |
| Right MFG | 447 | 3.6 | 34, 52, 18 | 4.2 |
| Left Cerebellum cortex | 193 | 1.5 | -30, -56, -30 | 4.3 |
| Left precentral gyrus | 147 | 1.2 | -40, -2, 52 | 3.2 |
| Right IFG | 141 | 1.1 | 42, 16, 22 | 3.2 |
| Left IFG | 109 | 0.9 | -40, 12, 28 | 3.1 |
| (b) Distributed > Selective |  |  |  |  |
| Region | # voxels | Size (cm<sup>3</sup>) | Peak (MNI coordinates) | t-statistics (sample mean) |
| Left STG | 2305 | 18.4 | -66, -18, 8 | 3.3 |
| Right STG | 1744 | 14.0 | 68, -18, 8 | 3.2 |
| (c) Selective > Distributed |  |  |  |  |
| Region | # voxels | Size (cm<sup>3</sup>) | Peak (MNI coordinates) | t-statistics (sample mean) |
| Right MFG | 250 | 2.0 | 38, 32, 20 | -2.8 |
| Left precentral gyrus | 218 | 1.7 | -28, -10, 52 | -2.6 |
| Left IPS | 190 | 1.6 | -26, -50, 46 | -3.4 |
| (d) Attention × Number of Speakers |  |  |  |  |
| Region | # voxels | Size (cm<sup>3</sup>) | Peak (MNI coordinates) | F-statistics (sample mean) |
| Left SMA | 367 | 3.0 | 10, 14, 50 | 7.0 |
| Right insula lobe | 320 | 2.6 | 34, 22, 6 | 7.2 |
| Left insula lobe | 240 | 2.0 | -34, 20, 2 | 6.8 |
| Right MFG | 193 | 1.6 | 32, 32, 26 | 7.3 |

STG, superior temporal sulcus; IFG, inferior frontal gyrus; MFG, middle frontal gyrus; IPS, inferior parietal sulcus; SMA, supplementary motor area.

**Figure 4.**
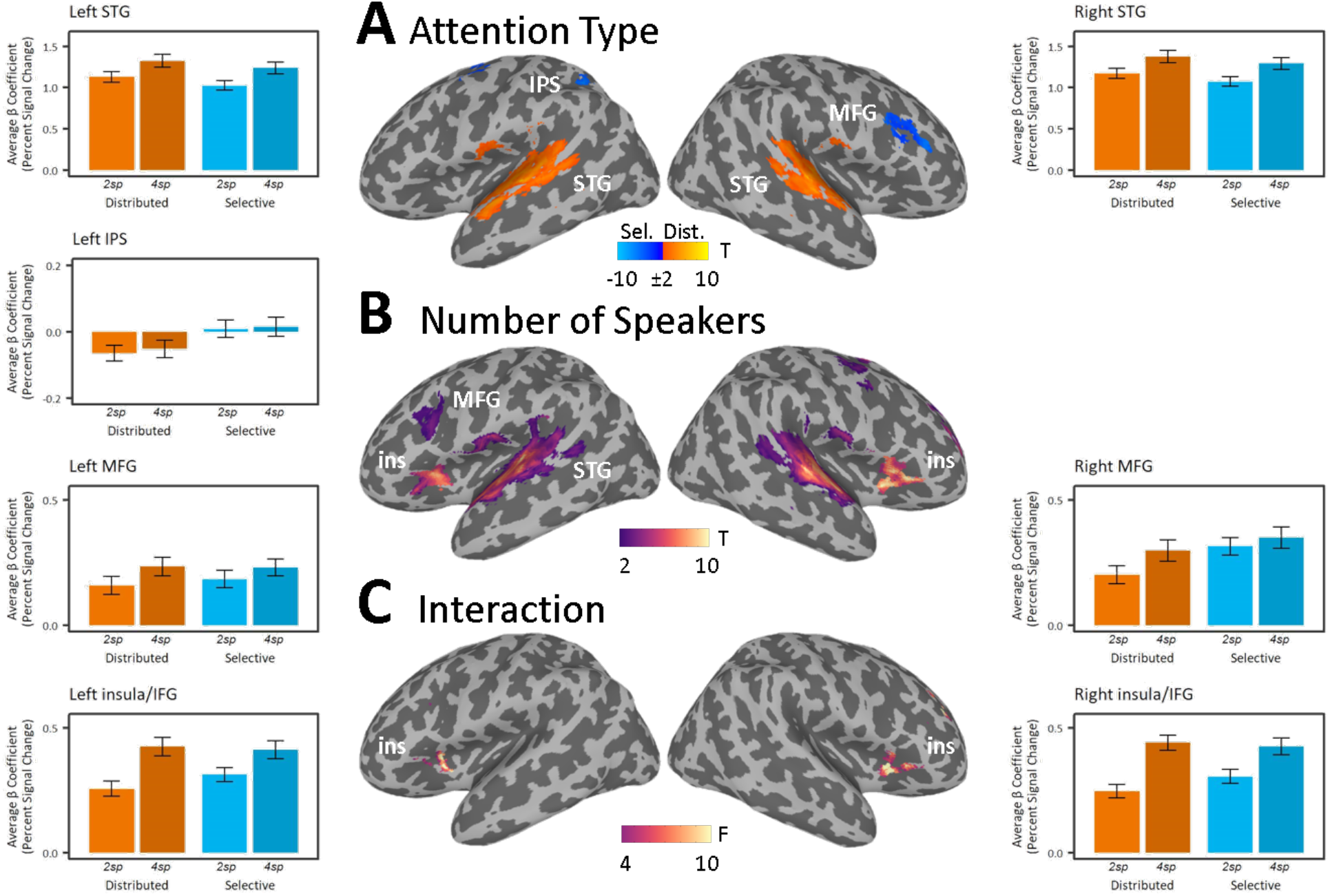
Results of a 2×2 ANOVA testing for effects of (A) Attention Type (Selective/Distrubuted), (B) Number of Speakers (2 / 4 speakers) and (C) their interaction (all maps thresholded at p<0.003, corrected). This analysis was performed within a mask of regions identified in Figure 3. Bar graphs on the two sides depict mean response and standard errors as a function of Attention Type (orange – Distributed, cyan – Selective) and Number of Speakers (2spk vs. 4spk) in each of the seven clusters with the most prominent statistical effects. Two areas (STG and IFG/insula) had where more than one statistically significant effect however within slightly different clusters. In these cases, data in the bar graph reflects the cluster exhibiting a main effect of Number of Speakers, however results are qualitatively similar if taken from the other significant cluster (for STG: main effect of Attention, for IFG/insula: the interaction; data not shown). IPS, inferior parietal sulcus; STG, superior temporal gyrus; MFG, middle frontal gyrus; ins, insula.

### 3.3. Functional connectivity results

Both types of attention engaged a widespread array of brain regions, including both sensory regions as well as fronto-parietal regions known to be engaged in speech processing and top-down attention control (Corbetta et al., 2008). In order to further ascertain the relationship between these regions, we computed the pairwise functional connectivity between them, and tested whether connectivity patterns were task-specific. This analysis focused on 7 ROIs where significant simple effects of Attention Type or Number of Speakers were found (see Table 5): bilateral auditory cortex, bilateral IPS, bilateral insula and right MFG. Functional connectivity was calculated between all ipsilateral ROI-pairs – yielding six pairs in the right hemisphere (R STG–MFG; R STG–IPS; R STG–insula; R MFG–IPS; R MFG–insula; R IPS–insula), and three in the left hemisphere (L STG–MFG; L STG–IPS; L STG–insula; Figure 5). Connectivity was calculated using the residual time-course of the BOLD signal between ROI-pairs, that captures the remaining variability in the data after accounting for main effects in the GLM. This was computed separately for each Attention Type and Number of Speakers (1, 2 & 4) as well as during the Rest periods.

**Figure 5.**
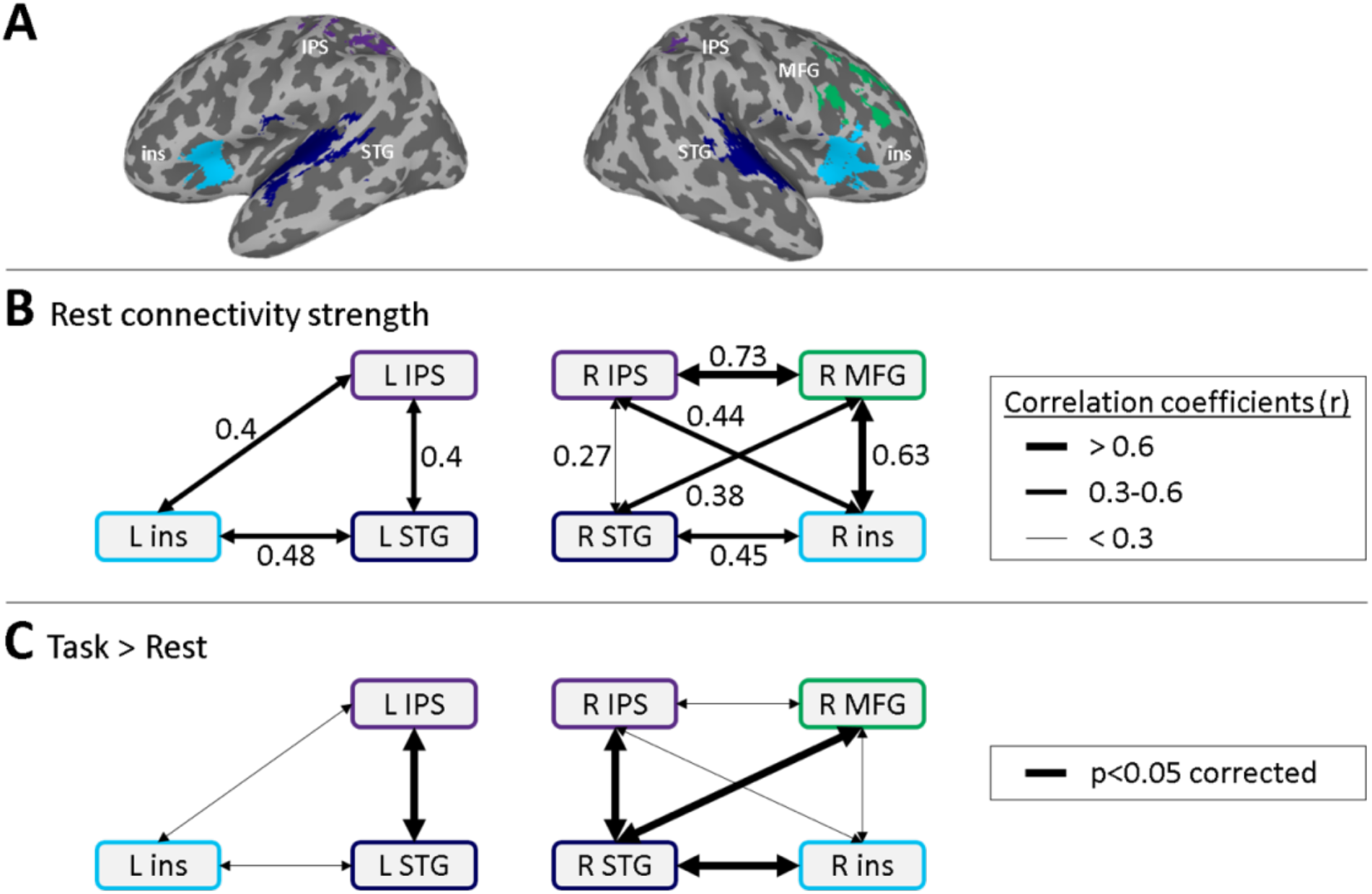
Functional connectivity results. (A) Depiction of the seven ROIs used for the connectivity analysis, derived from simple-effects analysis (see Methods). (B) Illustration of the connectivity strength between all ipsi-lateral ROI-pairs during the Rest periods. (C) Illustration of the ROI-pairs where connectivity strength was significantly increased during the experimental blocks (Task) vs. Rest. IPS, inferior parietal sulcus; STG, superior temporal gyrus; MFG, middle frontal gyrus; ins, insula.

Figure 5B illustrates the connectivity strengths between each ROI-pair during the Rest periods, when participants were not engaged in any task. Results show strong connectivity during Rest in the right hemisphere between the R-MFG and both the R-insula and the R-IPS. In line with previous work, this pattern suggests that these regions can be considered members of ‘default’ network connections (Van Calster et al., 2017; Alves et al., 2019). Next, we compared which regions became more strongly connected during the experimental conditions vs. Rest, which allowed us to discern task-related connectivity pattern and to test if these were modulated by the specific experimental condition. We found that four ROI-pairs, all involving auditory cortex in either hemisphere, showed significantly stronger connectivity during all experimental conditions vs. Rest (p<0.001, Bonferroni corrected for multiple comparisons): Three of them were in the right hemisphere (R STG–MFG, R STG–IPS, R STG–insula); and one in the left hemisphere (L STG– IPS; see Figure 5C). Importantly, however, connectivity strength was not significantly affected by the Attention Type required nor by the Number of Speakers in any of the 9 ROI-pairs tested (after Bonferroni correction). Table 6 provides a full report of the correlations between all ROI-pairs under the different Attention conditions, during the experimental and Rest blocks, with statistical analysis.

**Table 6.** Modulation of type of block (test/rest) on functional connectivity between ROIs

|  | Distributed Attention |  | Selective Attention |  | ANOVA results |  |  |
| --- | --- | --- | --- | --- | --- | --- | --- |
|  | Rest | Exp. | Rest | Exp. | Attention | Block | Interaction |
|  | r (SE) | r (SE) | r (SE) | r (SE) | F (p-value) | F (p-value) | F (p-value) |
| **(a) Left STG-insula** | 0.47 (0.03) | 0.52 (0.02) | 0.49 (0.03) | 0.52 (0.02) | 0.2 (0.6) | 4.6 (0.04) | 0.3 (0.6) |
| **(b) Left STG-IPS** | 0.39 (0.04) | 0.47 (0.03) | 0.41 (0.03) | 0.46 (0.03) | 0.01 (0.9) | **11 (0.002\*)** | 0.9 (0.35) |
| **(c) Left insula-IPS** | 0.38 (0.03) | 0.42 (0.03) | 0.42 (0.02) | 0.41 (0.03) | 0.6 (0.45) | 0.2 (0.15) | 0.3 (0.28) |
| **(d) Right STG-insula** | 0.43 (0.03) | 0.54 (0.02) | 0.47 (0.03) | 0.54 (0.02) | 1.7 (0.2) | **24.4**<br>**(<0.0001\*)** | 1.5 (0.2) |
| **(e) Right STG-IPS** | 0.24 (0.04) | 0.37 (0.03) | 0.3 (0.04) | 0.38 (0.03) | 2.3 (0.14) | **16.3**<br>**(0.003\*)** | 0.9 (0.4) |
| **(f) Right insula-IPS** | 0.44 (0.03) | 0.43 (0.03) | 0.45 (0.02) | 0.46 (0.03) | 0.5 (0.5) | 0.3 (0.6) | 0.6 (0.63) |
| **(g) Right MFG-STG** | 0.35 (0.04) | 0.49 (0.03) | 0.42 (0.03) | 0.48 (0.03) | 1.4 (0.3) | **22.2**<br>**(<0.0001\*)** | 6.5 (0.02) |
| **(h) Right MFG-IPS** | 0.73 (0.01) | 0.71 (0.02) | 0.73 (0.01) | 0.7 (0.02) | 0.001 (0.98) | 1.1 (0.3) | 0.3 (0.61) |
| **(i) Right MFG-insula** | 0.61 (0.02) | 0.62 (0.02) | 0.65 (0.02) | 0.64 (0.02) | 3.2 (0.09) | 1.6 (0.21) | 0.4 (0.54) |

Results of a 2×2 repeated-measures ANOVA (Selective/Distributed × rest/experimental). The attention conditions were collapsed across the 1- speaker, 2-speaker and 4-speaker blocks, since they did not exhibit any differences in functional connectivity. Dependent variable is the Z Fisher transformation of the r coefficients. Degrees of freedom for the F-values are all (1,31). The reported p-values are uncorrected; \*: significant p-values after a Bonferroni correction for multiple comparisons. Results in this table are qualitatively similar to the results of linear mixed effects regressions that include Attention (selective/distributed) and Number of Speakers (rest/1-sp/2-sp/4-sp) as fixed factors.

## 4. Discussion

Dealing effectively with input from concurrent speakers requires cooperation between the perceptual, linguistic and attentional systems. Here we found activation of a widespread and functionally connected array of brain regions, including bilateral auditory cortex, anterior-temporal language processing regions and some fronto-parietal regions, when participants heard input from multiple speakers relative to only a single speaker. Importantly, the same array of regions was activated regardless of what participants were instructed to ***do*** with the concurrent speech: ***attend selectively*** to a single speaker or ***distribute attention*** between all speakers. This pattern suggests that selective and distributed attention to speech engage a common neuroanatomical functional network (Lipschutz et al., 2002; Hill and Miller, 2010; Miller, 2015; Moisala et al., 2015). We did, however, find some difference in the magnitude of responses within these regions in the two attention conditions, which may reflect the different computations that each type of attention requires. Specifically, responses along bilateral STS/STG were enhanced during distributed attention, whereas responses in the right MFG and left IPS were enhanced when performing selective attention. The number of speakers also affected responses in several brain regions, most prominently along bilateral STS/STG and IFG/insulas, and in the latter there was also an interaction with Attention Type. In discussing the implications of these findings, it is worth reiterating which cognitive processes these two forms of attention have in common and the ways in which they differ.

Both selective and distributed attention to speech require perceptual analysis and segregation of a complex auditory scene into separate information streams, a process that becomes increasingly difficult as the number of concurrent speakers increases (Freyman et al., 2004; Simpson and Cooke, 2005). Hence, it makes sense that both types of attention generated activity in a large cluster along bilateral STG/STS, comprised of primary and associative auditory cortex, as well as more anterior portions involved in speech processing (identified independently as such in our functional speech localizer, and see: Humphries et al., 2001; Okada et al., 2010). Bilateral IFG/insula was also activated under both types of attention, consistent with its role in speech processing, particularly under adverse listening conditions or when speech is degraded (Davis and Johnsrude, 2003; Obleser et al., 2007; Davis et al., 2011; Alain et al., 2018). Increasing the number of competing speakers from two to four negatively impacted performance on both tasks. This also led to increase activity in both STG/STS and IFG/insula, for both types of attention, areas known to be sensitive to masking severity (Zatorre et al., 2004; Scott et al., 2009; Scott and McGettigan, 2013; Evans et al., 2016; Murphy et al., 2017; Peelle, 2018). This pattern indicates that increasing the perceptual load of the acoustic scene and, consequently, the difficultly to segregate competing streams, impacts both selective and distributed attention to speech.

However, selective and distributed attention differ in what the system ***does*** with the segregated streams. Distributed attention requires that all streams be fully segregated and processed – at an acoustic, phonetic, and semantic level. Hence, distributed attention to speech imposes a substantially higher perceptual and cognitive load on the listener than selective attention, where only a single stream needs to be fully processed (Gagné et al., 2017; Baldock et al., 2019; Lambez et al., 2020). Along these lines, the enhanced response observed in bilateral STG/STS for distributed vs. selective attention may reflect the number of auditory ‘objects’ that need to be represented. Indeed, previous dichotic listening paradigms, showed that responses increased in STG if both ears were to-be-attended vs. when the contralateral stimulus was to be selectively ignored (Hugdahl et al., 2000; Lipschutz et al., 2002). Similarly, we interpret the interaction found between Attention Type and Number of Speakers in bilateral IFG/insula as reflecting the increased challenge of maintaining linguist-level representations for multiple speech-streams during distributed attention.

Conversely, the main challenge of selective attention is enhancing the representation of a single task-relevant stream and relegating all other inputs to the perceptual-background to avoid distraction (Hillyard et al., 1973; Woldorff and Hillyard, 1991; Ghatan et al., 1998; Kawashima et al., 1999; Xiang et al., 2010; Ding and Simon, 2012). These selection and prioritization processes are carried out through top-down control mechanisms, involving fronto-parietal attention networks. This is in line with the enhanced responses found here during selective attention in two key members of fronto-parietal attention networks – right MFG and left IPS. Engagement of the left IPS is associated with voluntary top-down selection during both feature-based and spatial attention, in the auditory as well as the visual domain (Kastner and Ungerleider, 2001; Hill and Miller, 2010; Evans et al., 2016). Similarly, the right MFG is proposed to play a mediating role between the dorsal and ventral attention networks, controlling shifts of attention between top-down voluntary attention and reflexive ‘captures’ of attention (Fox et al., 2006; Corbetta et al., 2008; Petersen and Posner, 2012; Japee et al., 2015). Accordingly, the activation observed here in right MFG during selective attention might be attributed to avoiding unwanted attention shifts to irrelevant portion of the acoustic scene. This possibility is further supported by a previous study where right MFG responses to onsets of task-irrelevant speech were associated with their active rejection (Evans et al., 2016).

In thinking more broadly about the similarities and differences between selective and distributed attention to speech, it is important to recognize that these concepts can be operationalized in many different ways. In a recent study, similarly-motivated to ours, Yuriko Santos Kawata et al. (2020), measured the BOLD response to hearing two speakers simultaneously uttering different digits. They compared neural responses when participants knew in advance which speaker to respond to vs. when participants were required to decipher the digits spoken by both speakers. This variant of the selective vs. distributed attention manipulation imposed substantially higher acoustic and cognitive load than in the current study, due to the fully-overlapping speech-stimuli used and more complex stimulus-response mapping. Still, in both attention conditions they found activation of a network of temporo-fronto-parietal sensory, speech-processing and executive-control regions areas, that was very similar to one found here. Although in that study the effects of attention type within this network were somewhat different than those observed here, likely due to the vastly different stimuli used and task demands, the two studies converge in supporting a common temporo-fronto-parietal network underlying the allocation of attention to speech, with the ability to accommodate a variety of listening demands imposed in different conditions and contexts.

### Functional Connectivity

A complementary prism through which we studied the pattern of neural activations during selective and distributed attention to speech, was the functional connectivity between brain regions activated during this task. Connectivity analysis focused on the residual BOLD activity, i.e., fluctuations in the signal not explained by the GLM of task-related evoked response, which have been shown to carry task-relevant information independent of BOLD intensity (Rogers and Gore, 2008; Zhang and Li, 2010; Tran et al., 2018). This approach also allows inferring functional connectivity separately from the Rest and task blocks, providing an opportunity to determine which connectivity patterns reflect existing ‘default’, or ‘resting-state’ connections (Fair et al., 2007), and which are established dynamically for the purpose of executing a particular task. Here we found evidence for both types of connections. We observed high functional connectivity between MFG, IPS and the insula, particularly in the right hemisphere, which was similarly strong during Rest periods and task-blocks. This pattern is in line with their established membership in fronto-parietal ‘default’ attention networks (Fox and Raichle, 2007; Bullmore and Sporns, 2009). In contrast, bilateral STG was only weakly connected to the other regions during Rest, but its connectivity with almost all the other fronto-parietal ROIs increased significantly during the task. This pattern is indicative of task-specific ‘recruitment’ of sensory-regions into existing functional networks (Corbetta et al., 2008). Hence, the residuals-based connectivity analysis conducted here nicely demonstrates the two different types of network dynamics – endogenous default connections and transient task-specific connection – underscoring the system’s capability to carry out a diverse repertoire of perceptual and cognitive operations. Interestingly, we did not find significant differences in the connectivity patterns elicited under selective vs. distributed attention, nor any effects of Number of Speakers, a pattern that further supports the notion of a common underlying network for carrying out different attentional tasks and listening strategies.

## Conclusions

To summarize, attention to speech activates a large-scale network, comprised of brain regions involved in auditory processing, language-processing and top-down attentional control. The extensive overlap found here for selective and distributed attention, as well as the similar functional connectivity within this network during these distinct attention tasks, supports reliance on a common neural substrate. At the same time, the modulation of responses within these regions by the type of attention required and the number of competing speakers, reflects the flexibility of this network, its ability to implement different prioritization strategies among competing inputs and to adapt to different listener goals.

## Acknowledgements

This work was supported by the Israeli Ministry of Science and Technology [the Navon Fellowship (to GA)], the US-Israel Binational Science Foundation [grant number 2015385 (to EZG)], and the Israel Science Foundation [grant numbers 51/11 (to EZG, MBS) and 1083/17 (to MBS)].

## Notes

### Competing Interest Statement

The authors have declared no competing interest.

## References

Alain C, Du Y, Bernstein LJ, Barten T, Banai K (2018) Listening under difficult conditions: an activation likelihood estimation meta-analysis. Hum Brain Mapp 39:2695–2709.

Alves PN, Foulon C, Karolis V, Bzdok D, Margulies DS, Volle E, Thiebaut de Schotten M (2019) An improved neuroanatomical model of the default-mode network reconciles previous neuroimaging and neuropathological findings. Commun Biol 2:370.

Audacity (2018) Audacity ® | free, open source, cross-platform audio software for multi-track recording and editing. Audacity.

Baayen RH, Davidson DJ, Bates DM (2008) Mixed-effects modeling with crossed random effects for subjects and items. J Mem Lang 59:390–412.

Baldock J, Kapadia S, van Steenbrugge W (2019) The task-evoked pupil response in divided auditory attention tasks. J Am Acad Audiol 30:264–272.

Bates D, Maechler M, Bolker B, Walker S (2015) Fitting linear mixed-effects models using lme4. J Stat Softw 67:1– 48.

Broadbent DE (1958) Perception and Communication. London: Pergamon Press.

Bronkhorst AW (2015) The cocktail-party problem revisited: early processing and selection of multi-talker speech. Attention, Perception, Psychophys 77:1465–1487.

Bullmore E, Sporns O (2009) Complex brain networks: graph theoretical analysis of structural and functional systems. Nat Rev Neurosci 10:186–198.

Cooke M (2006) A glimpsing model of speech perception in noise. J Acoust Soc Am 119:1562–1573.

Corbetta M (1991) Selective and divided attention during visual discriminations of shape, color, and speed: functional anatomy by positron emission tomography. J Neurosci 11:2383–2402.

Corbetta M, Patel G, Shulman GL (2008) The reorienting system of the human brain: from environment to theory of mind. Neuron 58:306–324.

Cox RW (1996) AFNI: software for analysis and visualization of functional magnetic resonance neuroimages. Comput Biomed Res 29:162–173.

Davis MH, Ford MA, Kherif F, Johnsrude IS (2011) Does semantic context benefit speech understanding through “top-down” processes? evidence from time-resolved sparse fmri. J Cogn Neurosci 23:3914–3932.

Davis MH, Johnsrude IS (2003) Hierarchical processing in spoken language comprehension. J Neurosci 23:3423– 3431.

Ding N, Simon JZ (2012) Emergence of neural encoding of auditory objects while listening to competing speakers. Proc Natl Acad Sci 109:11854–11859.

Documentation M (2012) Matlab documentation. Matlab:R2012b.

Driver J (2001) A selective review of selective attention research from the past century. Br J Psychol 92 Part 1:53– 78.

Duncan J (1980) The locus of interference in the perception of simultaneous stimuli. Psychol Rev 87:272–300.

Evans S, Mcgettigan C, Agnew ZK, Rosen S, Scott SK (2016) Getting the cocktail party started: masking effects in speech perception. J Cogn Neurosci 28:483–500.

Fair DA, Schlaggar BL, Cohen AL, Miezin FM, Dosenbach NUF, Wenger KK, Fox MD, Snyder AZ, Raichle ME, Petersen SE (2007) A method for using blocked and event-related fmri data to study “resting state” functional connectivity. Neuroimage 35:396–405.

Fiedler L, Wöstmann M, Herbst SK, Obleser J (2019) Late cortical tracking of ignored speech facilitates neural selectivity in acoustically challenging conditions. Neuroimage 186:33–42.

Fox MD, Corbetta M, Snyder AZ, Vincent JL, Raichle ME (2006) Spontaneous neuronal activity distinguishes human dorsal and ventral attention systems. Proc Natl Acad Sci U S A 103:10046–10051.

Fox MD, Raichle ME (2007) Spontaneous fluctuations in brain activity observed with functional magnetic resonance imaging. Nat Rev Neurosci 8:700–711.

Freyman RL, Balakrishnan U, Helfer KS (2004) Effect of number of masking talkers and auditory priming on informational masking in speech recognition. J Acoust Soc Am 115:2246–2256.

Fritz JB, Elhilali M, David S V., Shamma SA (2007) Auditory attention - focusing the searchlight on sound. Curr Opin Neurobiol 17:437–455.

Gagné JP, Besser J, Lemke U (2017) Behavioral assessment of listening effort using a dual-task paradigm: a review. Trends Hear 21:1–25.

Getzmann S, Golob EJ, Wascher E (2016) Focused and divided attention in a simulated cocktail-party situation: erp evidence from younger and older adults. Neurobiol Aging 41:138–149.

Ghatan PH, Hsieh JC, Petersson KM, Stone-Elander S, Ingvar M (1998) Coexistence of attention-based facilitation and inhibition in the human cortex. Neuroimage 7:23–29.

Gygi B, Shafiro V (2012) Spatial and temporal factors in a multitalker dual listening task. Acta Acust united with Acust 98:142–157.

Haegens S, Zion Golumbic EM (2018) Rhythmic facilitation of sensory processing: a critical review. Neurosci Biobehav Rev 86.

Hambrook DA, Tata MS (2019) The effects of distractor set-size on neural tracking of attended speech. Brain Lang 190:1–9.

Hill KT, Miller LM (2010) Auditory attentional control and selection during cocktail party listening. Cereb Cortex 20:583–590.

Hillyard S, Hink R, Schwent V, Picton T (1973) Electrical signs of selective attention in the human brain. Science (80-) 182:177–180.

Hugdahl K, Law I, Kyllingsbæk, Brønnick K, Gade A, Paulson OB (2000) Effects of attention on dichotic listening: an 15o-pet study. Hum Brain Mapp 10:87–97.

Humphries C, Willard K, Buchsbaum B, Hickok GCA (2001) Role of anterior temporal cortex in auditory sentence comprehension: an fmri study. Neuroreport 12:1749–1752.

Jaeger TF (2008) Categorical data analysis: away from anovas (transformation or not) and towards logit mixed models. J Mem Lang 59:434–446.

Jäncke L, Shah NJ (2002) Does dichotic listening probe temporal lobe functions? Neurology 58:736–743.

Japee S, Holiday K, Satyshur MD, Mukai I, Ungerleider LG (2015) A role of right middle frontal gyrus in reorienting of attention: a case study. Front Syst Neurosci 9:23.

Johannsen P, Jakobsen J, Bruhn P, Hansen SB, Gee A, Stødkilde-Jørgensen H, Gjedde A (1997) Cortical sites of sustained and divided attention in normal elderly humans. Neuroimage 6:145–155.

Johnson JA, Zatorre RJ (2006) Neural substrates for dividing and focusing attention between simultaneous auditory and visual events. Neuroimage 31:1673–1681.

Jones MR (2019) Time will tell: A theory of dynamic attending. Oxford University Press.

Kastner S, Ungerleider LG (2001) The neural basis of biased competition in human visual cortex. Neuropsychologia 39:1263–1276.

Kawashima R, Imaizumi S, Mori K, Okada K, Goto R, Kiritani S, Ogawa A, Fukuda H (1999) Selective visual and auditory attention toward utterances - a pet study. Neuroimage 10:209–215.

Kawashima T, Sato T (2015) Perceptual limits in a simulated “cocktail party.” Attention, Perception, Psychophys 77:2108–2120.

Koelewijn T, Shinn-Cunningham BG, Zekveld AA, Kramer SE (2014) The pupil response is sensitive to divided attention during speech processing. Hear Res 312:114–120.

Kuznetsova A, Brockhoff PB, Christensen RHB (2017) LmerTest package: tests in linear mixed effects models. J Stat Softw 82.

Lambez B, Agmon G, Har-Shai P, Rassovsky Y, Zion-Golumbic EM (2020) Paying attention to speech: the role of cognitive capacity and acquired experience. Attention, Percept Psychophys.

Lavie N, Beck DM, Konstantinou N (2014) Blinded by the load: attention, awareness and the role of perceptual load. Philos Trans R Soc Lond B Biol Sci 369:20130205.

Lavie N, Hirst A, de Fockert JW, Viding E (2004) Load theory of selective attention and cognitive control. J Exp Psychol Gen 133:339–354.

Lipschutz B, Kolinsky R, Damhaut P, Wikler D, Goldman S (2002) Attention-dependent changes of activation and connectivity in dichotic listening. Neuroimage 17:643–656.

Loose R, Kaufmann C, Auer DP, Lange KW (2003) Human prefrontal and sensory cortical activity during divided attention tasks. Hum Brain Mapp 18:249–259.

McCloy DR, Lee AKC (2015) Auditory attention strategy depends on target linguistic properties and spatial configuration. J Acoust Soc Am 138:97–114.

Miller LM (2015) Neural mechanisms of attention to speech. In: Neurobiology of Language, pp 503–514. Elsevier Inc.

Moisala M, Salmela V, Salo E, Carlson S, Vuontela V, Salonen O, Alho K (2015) Brain activity during divided and selective attention to auditory and visual sentence comprehension tasks. Front Hum Neurosci 9:86.

Murphy S, Spence C, Dalton P (2017) Auditory perceptual load: a review. Hear Res 352:40–48.

Obleser J, Wise RJS, Dresner MA, Scott SK (2007) Functional integration across brain regions improves speech perception under adverse listening conditions. J Neurosci 27:2283–2289.

Okada K, Rong F, Venezia J, Matchin W, Hsieh IH, Saberi K, Serences JT, Hickok G (2010) Hierarchical organization of human auditory cortex: evidence from acoustic invariance in the response to intelligible speech. Cereb Cortex 20:2486–2495.

Peelle JE (2018) Listening effort: how the cognitive consequences of acoustic challenge are reflected in brain and behavior. Ear Hear 39:204–214.

Petersen SE, Posner MI (2012) The attention system of the human brain: 20 years after. Annu Rev Neurosci:73–89.

Posner MI (1980) Orienting of attention. Q J Exp Psychol:3–25.

Rogers BP, Gore JC (2008) Empirical comparison of sources of variation for fmri connectivity analysis Heo M, ed. PLoS One 3:e3708.

Satterthwaite FE (1946) An approximate distribution of estimates of variance components. Biometrics Bull 2:110.

Scott SK, McGettigan C (2013) The neural processing of masked speech. Hear Res 303:58–66.

Scott SK, Rosen S, Beaman CP, Davis JP, Wise RJS (2009) The neural processing of masked speech: evidence for different mechanisms in the left and right temporal lobes. J Acoust Soc Am 125:1737–1743.

Simpson SA, Cooke M (2005) Consonant identification in n-talker babble is a nonmonotonic function of n. J Acoust Soc Am 118:2775–2778.

Stoppelman N, Harpaz T, Ben-Shachar M (2013) Do not throw out the baby with the bath water: choosing an effective baseline for a functional localizer of speech processing. Brain Behav 3:211–222.

Taylor PA, Saad ZS (2013) FATCAT: (an efficient) functional and tractographic connectivity analysis toolbox. Brain Connect 3:523–535.

Tóth B, Farkas D, Urbán G, Szalárdy O, Orosz G, Hunyadi L, Hajdu B, Kovács A, Szabó BT, Shestopalova LB, Winkler I (2019) Attention and speech-processing related functional brain networks activated in a multi-speaker environment. PLoS One 14:e0212754.

Tran SM, McGregor KM, James GA, Gopinath K, Krishnamurthy V, Krishnamurthy LC, Crosson B (2018) Task-residual functional connectivity of language and attention networks. Brain Cogn 122:52–58.

Treisman AM (1964) The effect of irrelevant material on the efficiency of selective listening. Am J Psychol 77:533– 546.

Van Calster L, D’Argembeau A, Salmon E, Peters F, Majerus S (2017) Fluctuations of attentional networks and default mode network during the resting state reflect variations in cognitive states: evidence from a novel resting-state experience sampling method. J Cogn Neurosci 29:95–113.

Vestergaard MD, Fyson NRC, Patterson RD (2011) The mutual roles of temporal glimpsing and vocal characteristics in cocktail-party listening. J Acoust Soc Am 130:429–439.

Woldorff MG, Gallen CC, Hampson SA, Hillyard SA, Pantev C, Sobel D, Bloom FE (1993) Modulation of early sensory processing in human auditory cortex during auditory selective attention. Proc Natl Acad Sci U S A 90:8722–8726.

Woldorff MG, Hillyard SA (1991) Modulation of early auditory processing during selective listening to rapidly presented tones. Electroencephalogr Clin Neurophysiol 79:170–191.

Xiang J, Simon J, Elhilali M (2010) Competing streams at the cocktail party: exploring the mechanisms of attention and temporal integration. J Neurosci 30:12084–12093.

Yost WA, Dye RH, Sheft S (1996) A simulated “cocktail party” with up to three sound sources. Percept Psychophys 58:1026–1036.

Yuriko Santos Kawata N, Hashimoto T, Kawashima R (2020) Neural mechanisms underlying concurrent listening of simultaneous speech. Brain Res 1738:146821.

Zatorre RJ, Bouffard M, Belin P (2004) Sensitivity to auditory object features in human temporal neocortex. J Neurosci 24:3637–3642.

Zhang S, Li CR (2010) A neural measure of behavioral engagement: task-residual low-frequency blood oxygenation level-dependent activity in the precuneus. Neuroimage 49:1911–1918.

Zion Golumbic EM, Ding N, Bickel S, Lakatos P, Schevon CA, McKhann G, Goodman RR, Emerson R, Mehta AD, Simon JZ, Poeppel D, Schroeder CE (2013) Mechanisms underlying selective neuronal tracking of attended speech at a cocktail party. Neuron 77:980–991.

Zion Golumbic EM, Poeppel D, Schroeder CE (2012) Temporal context in speech processing and attentional stream selection: a behavioral and neural perspective. Brain Lang 122.

